# Understanding How Microbiomes Influence The Systems They Inhabit: Moving From A Correlative To A Causal Research Framework

**DOI:** 10.1101/065128

**Authors:** E.K. Hall, E.S Bernhardt, R.L. Bier, M.A. Bradford, C.M. Boot, J.B. Cotner, P.A. del Giorgio, S.E. Evans, E.B. Graham, S.E. Jones, J.T. Lennon, K.J. Locey, D. Nemergut, B.B. Osborne, J.D. Rocca, J.S. Schimel, M.P. Waldrop, M.W. Wallenstein

## Abstract

Translating the ever-increasing wealth of information on microbiomes (environment, host, or built environment) to advance the understanding of system-level processes is proving to be an exceptional research challenge. One reason for this challenge is that relationships between characteristics of microbiomes and the system-level processes they influence are often evaluated in the absence of a robust conceptual framework and reported without elucidating the underlying causal mechanisms. The reliance on correlative approaches limits the potential to expand the inference of a single relationship to additional systems and advance the field. We propose that research focused on how microbiomes influence the systems they inhabit should work within a common framework and target known microbial processes that contribute to the system-level processes of interest. Here we identify three distinct categories of microbiome characteristics (microbial processes, microbial community properties, and microbial membership) and propose a framework to empirically link each of these categories to each other and the broader system level processes they affect. We posit that it is particularly important to distinguish microbial community properties that can be predicted from constituent taxa (community aggregated traits) from and those properties that are currently unable to be predicted from constituent taxa (emergent properties). Existing methods in microbial ecology can be applied to more explicitly elucidate properties within each of these categories and connect these three categories of microbial characteristics with each other. We view this proposed framework, gleaned from a breadth of research on environmental microbiomes and ecosystem processes, as a promising pathway with the potential to advance discovery and understanding across a broad range of microbiome science.

## Current Approaches Linking Microbial Characteristics and Ecosystem Processes

Virtually all ecosystem processes are influenced by microorganisms, and many processes are carried out exclusively by microorganisms. This has sometimes led to the assumption that a better description of the microbiome (including its associated transcripts, proteins, and metabolic products) should lead to a better understanding and predictions of system level processes. However, such justifications assume that measurable characteristics of the microbiome (e.g. 16S rRNA gene libraries, metagenomes, enzymatic activities) can inform our ability to better understand and predict system-level processes. Unfortunately, additional information about the microbiome does not always provide a clearer understanding of ecosystem processes beyond what can be predicted by environmental factors alone^1,2^.

Two recent meta-analyses^3,4^ suggest that research at the intersection of ecosystem science and microbial ecology often relies on assumed relationships between microbiome characteristics and ecosystem processes and often do not test to see if those relationships are present. The first, an examination of 415 studies, found little evidence that protein-encoding genes (sometimes referred to as “functional genes”) or gene transcripts correlate with associated biogeochemical processes^3^. Although all studies attempted (or presumed) to link microbial genes or transcripts with function, only 14% measured both the abundance of genes or transcripts and the corresponding process. When the relationship between microbial characteristic and ecosystem process was measured and tested only 38% exhibited a positive correlation^3^. This result was consistent whether functional gene or transcript abundance was used as the response variable.

The second study compiled a separate dataset of 148 studies that examined microbial membership and ecosystem processes in response to experimental manipulations^4^. Whereas 40% of included studies reported concomitant changes in microbial membership and an ecosystem process, only one third of those cases reported the relationship between microbial membership and an ecosystem process. Interestingly, of the 53 studies that posed a hypothesis about links between microbial membership and ecosystem processes, more than half (53%) did not report testing for a statistical link between membership and processes^4^.

Microbiomes are the engines that power system-level processes^5^. However the meta-analyses described above illustrate that the current approach to study the links between microbiome characteristics and ecosystem processes are not well formulated and relationships between microbiome characteristics and system level processes are implied yet rarely tested. When linkages are tested, significant correlations between microbiome characteristics and ecosystem processes are sometimes present, but more frequently not^3,4^. One reason for the ambiguity between microbiome characteristics and system level processes is that many studies are conducted in the absence of a conceptual framework that illustrates how different measurable microbial characteristics relate to one another and to the system level process of interest.

## Challenges in Linking Microbial Characteristics and Ecosystem Processes

Current conceptual research frameworks that attempt to link microbial information to a system-level process often do not effectively align with the methods being applied or the data those methods generate. For example, environmental factors act on the physiology of individual organisms, which alters their competitive ability, abundance, and ultimately their contribution to an ecosystem process. However, designing an observational study or experiment from this framework assumes that environmental and microbial characteristics are measurable across multiple categories of ecological organization (i.e., individuals, populations, and communities) at the temporal and spatial scales at which they influence system level processes (Figure 1a). In addition, the relationships between environmental variables and microbial characteristics can be decoupled in both time and space^4^, and are often non-linear^6^. Recent immigration, phenotypic plasticity, disequilibrium between the environment and the extant microbiome at the time of sampling, functional redundancy, and dormancy can all mask the relationship between measurable microbial characteristics and the processes microorganisms influence (Figure 1b).^4,7,8,9^ As micrometer scale characteristics of microbiomes (10^−6^ m) are scaled to the level of ecosystems (m to km), we assume that our conceptual understanding is also scalable. However, each of the aforementioned confounding factors aggregate over multiple orders of magnitude often masking the very relationships we seek to elucidate (Figure 1b). To formalize how measurable microbiome characteristics are linked with system-level processes we have conceptually defined the intersection of microbial and ecosystem ecology and identified three categories of microbial characteristics to illustrate how they interact with each other and may contribute to an ecosystem process (Figure 2).

**Figure 1.**
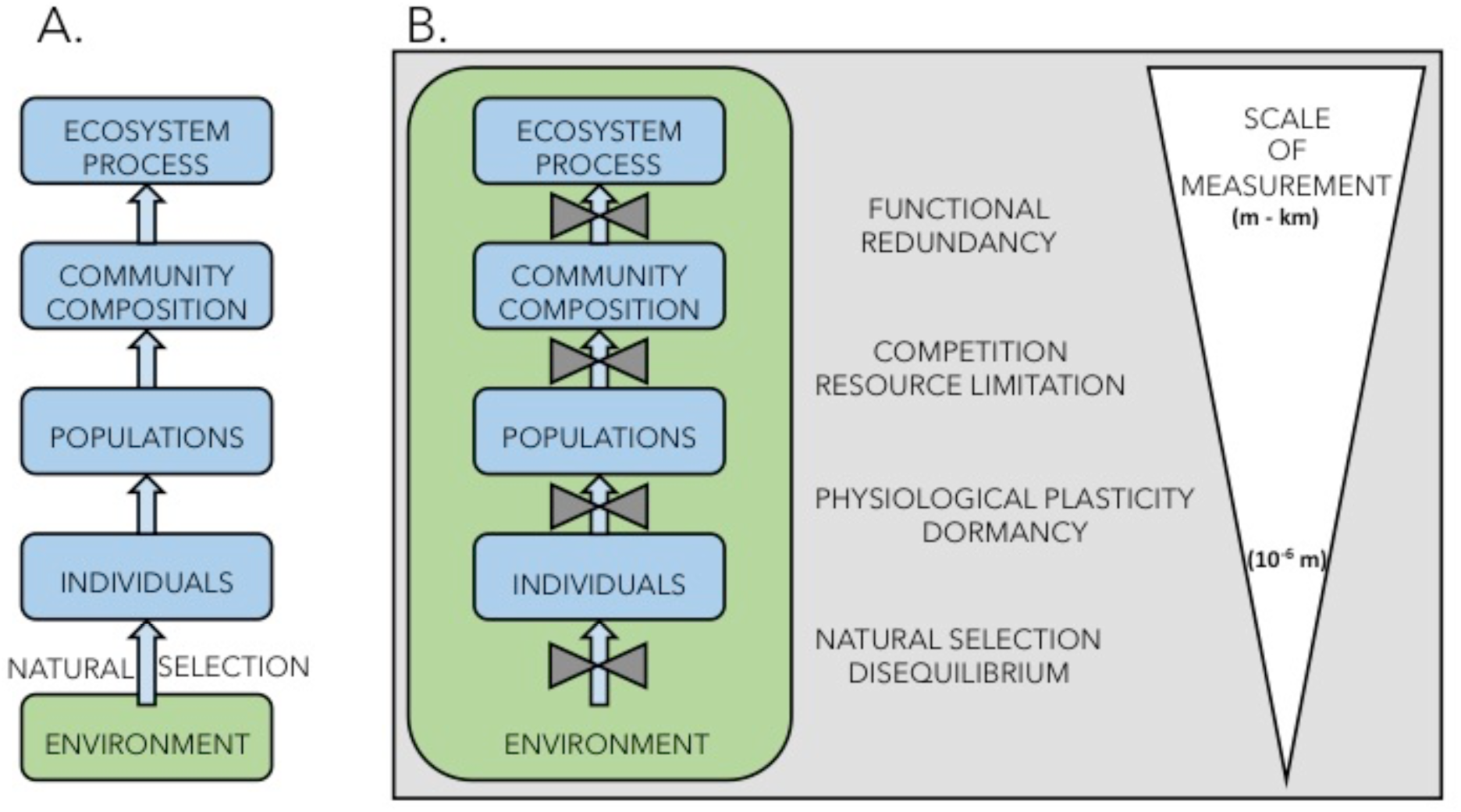
Diagram of microbial-ecosystem linkages A) how linkages are commonly conceptualized across levels of ecological organization and B) the series of ecological phenomena that create challenges when attempting to link metrics from one level of ecological organization to the other.

**Figure 2.**
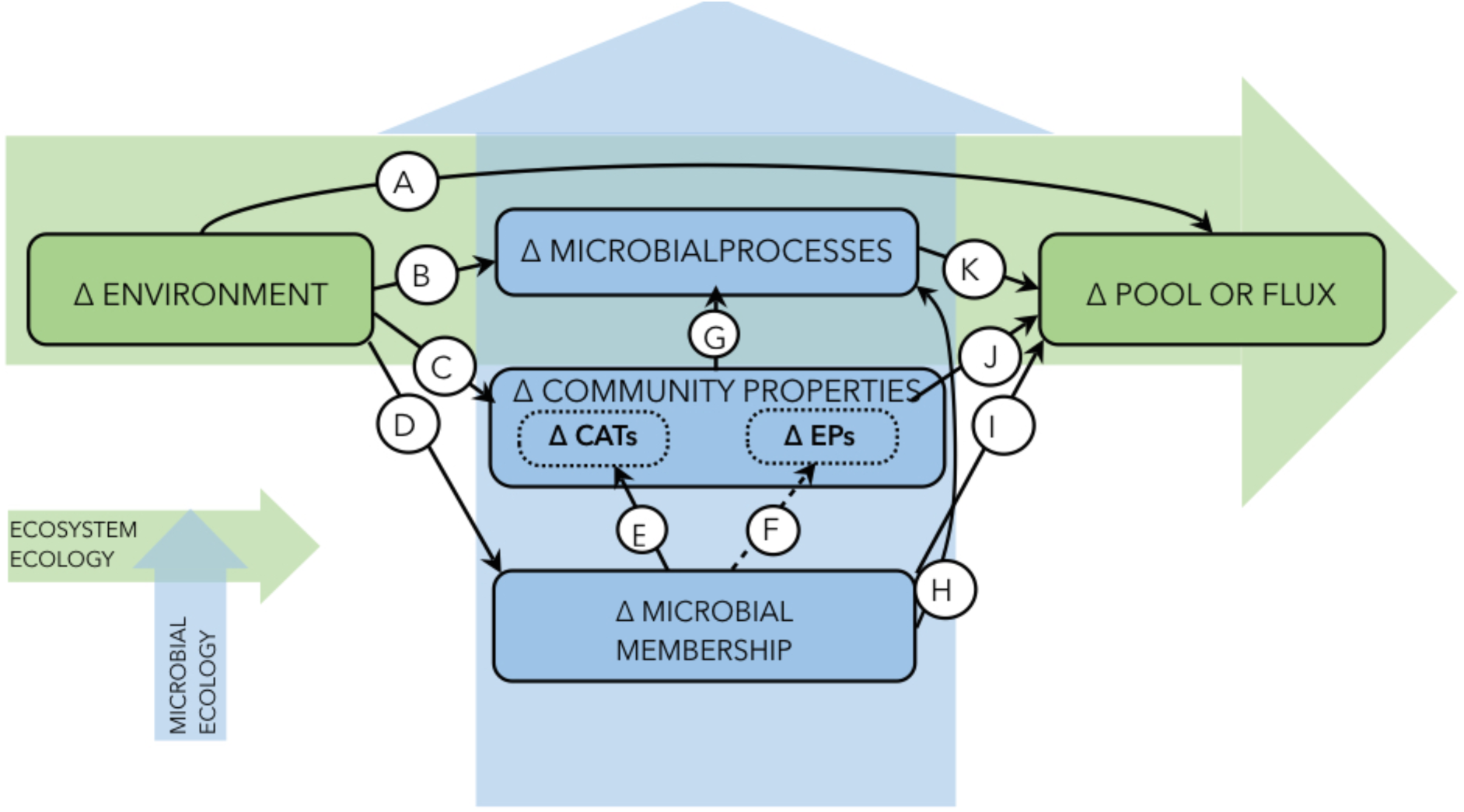
A conceptual map of the intersection between microbial (vertical) and ecosystem (horizontal) ecology illustrating each of the three categories of microbial characteristics (microbial processes, community properties, and microbial membership) as defined in the text. We argue for an increased focus on studies that elucidate pathways E, F, and G. In addition, we note that pathways I and J are less likely to effectively incorporate microbiome characteristics into system-level science. The delta symbol in each category indicates an emphasis on how changes within a category may lead to a change in a connected category. The dotted arrow for letter F denotes that many emergent properties cannot currently be linked to membership and is an important area for active research. CAT = community aggregated trait, EP=emergent property.

## Mapping Ecosystem Processes to Microbial Characteristics

Defining the ecosystem process, its critical sub-processes, and the known phylogenetic distribution of the metabolic pathways that drive those sub-processes creates an explicit conceptual pathway that links the ecosystem process to the microorganisms that contribute to it. Ecosystem processes are defined as qualitative changes in a pool, or a flux from one pool to another (e.g., NH_4_^+^ to NO_3_^−^, or dissolved organic matter to CO_2_). The first step to understand how a microbiome influences an ecosystem process is to define the ecosystem process of interest and each sub-process that contributes to it (i.e., the set of constituent reactions that combine to determine a net flux). Ecosystem processes are composites of complementary or antagonistic sub-processes, carried out by phylogenetically and metabolically diverse microorganisms^10^. For example, net ecosystem productivity (NEP) is the balance between antagonistic processes of C-fixation and C-mineralization. Each sub-process of NEP can be further partitioned into a series of metabolic pathways (e.g., chemoautotrophic nitrification and photoautotrophic C-fixation or heterotrophic fermentation and aerobic respiration). Partitioning each ecosystem process in this hierarchical manner can continue until the sub-processes maps directly to specific microbial metabolic pathways (e.g., acetoclastic methanogenesis). Subsequently, each of these metabolic pathways can be categorized as either phylogenetically broad or narrow^11^. Broad processes are phylogenetically common (i.e., widely distributed among taxa), whereas narrow processes are phylogenetically conserved (i.e., limited to a specific subset of taxa). For example, denitrification and photosynthesis are phylogenetically broad processes, while both methanogenesis and methanotrophy are currently thought to be phylogenetically narrow processes (with at least one notable exception^12^).

The second step is to identify the controls or constraints on each constituent sub-process. For example, the kinetics of a single metabolic pathway in a model organism may help us understand the rate limiting steps of a narrow process, but insights from model organisms are much less likely to capture the full spectrum of responses of a broad process where phenotypic variation among phylogenetically diverse organisms is likely to be much greater^13,14^. Once the ecosystem process has been conceptually partitioned into its component parts and their primary controls, a concerted approach can be applied to investigate how characteristics of the microbiome influence the ecosystem process of interest within the complexity of a natural environment.

## Categories of Microbial Characteristics

The contribution of the microbiome to ecosystem processes is exerted through aggregate community properties that are shaped by both microbial membership and environmental factors. Most current studies rarely articulate how measurements of microbial characteristics differ in their specificity (i.e., the level of phylogenetic resolution), precision (i.e., the ability of the method to repeatedly describe the characteristic of interest), or context (i.e., how a characteristic relates to other microbial characteristics or the ecosystem). To aid in designing experiments that link microorganisms to the processes they influence we propose three distinct categories of microbial characteristics: 1) microbial processes, 2) microbial community properties, and 3) microbial membership (Figure 2). This proposed framework allows the researcher to clearly identify how different measurements used to characterize a microbiome interact with each other, and to identify the potential of each characteristic to elucidate the microbial contribution to the system level process. All measurable characteristics of microbial communities (e.g., abundance of cells, sequence of genes, transcripts, or proteins; enzyme expression or activity) can be placed within one of the above categories. This conceptual structure that orientates each microbial category within a broader context creates the opportunity to improve the design of observational and experimental studies in microbiome research. *Microbial Processes* - Microbial processes are the collective metabolisms of the microbiome that contribute to changes in pools and fluxes of elements or compounds (i.e., Figure 2, Letter K). This is the category of microbial information that can most readily be incorporated into system-level models because many microbial processes represent the key sub-processes that contribute to a particular ecosystem process (e.g., methanogenesis + methanotrophy ≈ methane efflux). Commonly measured microbial processes in ecosystem science include nitrogen fixation, denitrification, nitrification, phosphorus uptake and immobilization, carbon fixation, and organic carbon mineralization. The rates of many microbial processes can be approximated through physiological assays (e.g., biological oxygen demand to estimate microbial respiration), and while they do not open the “black box” of the microbial community, they do directly quantify the microbial contribution (or potential contribution) to changes in resources moving through the box. Microbial processes can be distinguished from other microbial characteristics because they are all rates (i.e., have time in the denominator) and require a bioassay to estimate.

Assays used to estimate microbial processes are often logistically challenging, require manipulations that inevitably deviate from *in situ* conditions, and often depend on the environment from which the microbiome was sampled. For example, the relationship between temperature and microbial processes such as enzyme activity and phosphorus use efficiency vary across latitudinal gradients^15^ and among seasons^16^, respectively. Thus, observations of the effect of temperature on either enzyme activity or phosphorus use efficiency depend on where (e.g., at what latitude) and when (e.g., during which season) they were measured. In the absence of an understanding of the underlying physiological mechanism (e.g., the physiological change that allows a community to perform differently at different temperatures), the relationship between and environmental driver (e.g., temperature) and a microbial process must be measured through a direct assay at each location and at each time. This limits the inference possible from relying on measurements of microbial processes alone to understand the microbial contribution to an ecosystem process.

*Microbial Community Properties* - Microbial community properties include a broad set of microbial characteristics such as community biomass or biomass elemental ratios (e.g., biomass C:N or C:P ratios) and the majority of phylogenetically undifferentiated aggregate sequence based measurements (e.g., gene abundance, metagenomes, transcriptomes). Microbial community properties (Figure 2) represent an integrated characteristic of the microbiome that has the potential to predict or at least constrain the estimates of microbial processes. For example, microbial community biomass C:N (a community property) has been shown to indicate a microbiome’s potential to mineralize or immobilize N in terrestrial^17^, freshwater^18^ and marine^19^ ecosystems.

Microbial community properties can be separated into two categories, emergent properties and community aggregated traits. It is generally agreed that emergent properties refer to a quality of the whole that is unique and distinguishable from the additive properties of its constituents. Whereas in some cases, emergent properties may be predicted from their constituent parts, for example, prediction of the physical properties of carbon polymers is possible based on the atomic organization of carbon atoms within each structure^20^, in microbial ecology our understanding of the constituent parts and their interactions are more often than not insufficient to predict emergent properties. Here we use emergent properties as it has been defined previously in ecology^21^: “An emergent property of an ecological unit is one which is wholly unpredictable from observation of the components of that unit”, which is also consistent with its contemporary use in microbial ecology_^22^_.

Emergent properties of microbiomes influence important ecosystem processes. For example, a series of experiments in flow-through flumes mimicked development and metabolism of stream biofilms^23^. Transient storage (i.e., an increase in residence time of the water and its solutes near the biofilm relative to the flow around it) increased as the microbial biofilm density increased.^23^ In this case biofilm density was an emergent property of the microbiome and transient storage was the microbial process it mediated. Another example of an emergent property is the distribution of traits that influences key microbial processes. Trait based approaches have a rich history in ecology and have been increasingly applied to address questions in multiple areas of microbial ecology.^24^ For example, uptake of an organic substrate can often involve the expression of multiple genes, differing among individual organisms, all capable of performing uptake of the organic substrate, albeit with differences in the underlying efficiency. The distribution of the expression of these functional gene variants generates a trait distribution that is an emergent property of the microbiome. That emergent trait distribution determines the overall performance of the microbiome for that function (i.e., the uptake of the organic substrate), but it cannot be predicted from the presence of the organisms conferring that trait using current methods.^25^ While characterization of emergent properties may improve the understanding of microbial processes (Figure 2, Letter G) currently they most often cannot be estimated or predicted on the basis of the constituent taxa (i.e. membership) alone (Figure 2, Letter F), and thus must remain as an intermediary between environmental drivers (Figure 2, Letter C) and microbial processes (Figure 2, Letter G).

Unlike emergent properties, community aggregated traits can potentially be estimated from characteristics of their constituents and provide a pathway to link microbial community membership to the community properties that drive microbial processes (Figure 2, Letter E).^26^ For example, community aggregated traits may include commonly measured community properties such as functional gene abundance as estimated from qPCR (e.g., *pmo*A which encodes a subunit of the enzyme involved in methane oxidation, can be used to estimate potential for methanotrophy and as a phylogenetic marker for methanotrophs)^27^. A recent perspective article that discussed the role of community aggregated traits in microbial ecology noted a series of additional putative community aggregated traits (e.g., maximum growth rate, dormancy, osmoregulation) that could be inferred from metagenomic data of the extant community.^26^

Understanding which community properties can be predicted by membership is a critical research question and an important step in understanding how the microbiome contributes to system level processes. Whether or not a community property is likely to be an emergent property or a community aggregated trait is an exciting area of research and provides an important framework to advance research at the microbial-ecosystem nexus. New approaches, like studying higher-level interactions in ecological communities could help understand how a microbiome’s constituents interact to from emergent properties.^28^ This is not a trivial task, yet a suite of existing methods, discussed below, provides the ability to directly pursue this challenge.

*Microbial Community Membership* **-** Although the now commonplace analysis of community membership by sequencing phylogenetic markers (e.g., ITS regions or regions of the16S rRNA and 18S rRNA genes) or suites of phylogenetically conserved protein sequences identifies constituent microbial taxa, the direct coupling of microbial phylogeny to physiology and ecology remains elusive (Figure 2, Letter H).^29,30,31^ In general the paucity of associated physiological data or information on population phenotypes that accompany phylogenetic analyses limits the system-level inference that is possible from analyses of community membership. Even when the physiology of an organism is known, it appears many metabolic pathways are phylogenetically broad, and that any given microbiome will contain a diverse set of microorganisms with the genes that encode many of the same common microbial metabolic pathways, often referred to as “functional redundancy”^32^. There also appears to be no consistent phylogenetic resolution at which specific microbial metabolic pathways are constrained^32^. This constrains our ability to attribute microbial processes to community membership of even relatively simple environmental consortia. Whereas it is clear that microbial populations are not randomly distributed in space and time^31^, and that some microbial traits are conserved at coarse taxonomic scales, ^24,33,34^ the physiological mechanisms underlying non-random distributions of microbial taxa across environmental gradients are often unknown. This limited understanding of the metabolism of most bacterial phyla is one thing that currently prevents linking a microorganism’s abundance in an environment to its role in a related microbial process.

## A Path Forward

We suggest that a challenging but necessary step for microbiome science is to move away from identifying correlative relationships between characteristics of the microbiome and system level processes, and towards identifying more causative and mechanistic relationships. The conceptual diagram (Figure 2) is a road map to organize and link the diverse suite of measurable microbial characteristics that are currently available to researchers. Figure 2 does not represent how these components necessarily interact in the environment; rather it is a map that identifies potential links between measurable microbial and system-level characteristics that can help structure our exploration of how microorganisms influence the systems they inhabit. Ecosystem ecology has traditionally been confined to interactions between environmental parameters and ecosystem processes (depicted within the horizontal arrow, Figure 2). Similarly, microbial ecology (depicted within the vertical arrow, Figure 2) has historically focused on phylogenetically undefined aspects of microbial communities (e.g., bacterial abundance) and microbial processes (e.g., bacterial production), or the physiology of microbial isolates (e.g., sulfate reducing bacteria) or the collective physiology of highly reduced communities with known membership (e.g., waste water treatment microbiome). The routine inclusion of sequence-based approaches in studies of environmental microorganisms has led to an increasingly detailed description of the world’s microbiomes and an increasing interest in how constituents of those microbiomes interact to influence the system as a whole.

Currently, researchers use a range of approaches for attempting to link characteristics of the microbiome to ecosystem processes. Direct connections between microbial membership and ecosystem processes (Figure 2, Letter I), or community properties and ecosystem processes (Figure 2, Letter J), have proven difficult to establish.^3,4^ We propose 1) identifying which microbial processes are likely to contribute to ecosystem-level pools and fluxes *a priori* (Figure 2, Letter K), 2) determining which microbial community properties best describe and predict these microbial processes (Figure 2, Letter G), and 3) identifying whether the community properties that best describe each process are a community aggregated trait or a emergent property (Community Properties, Figure 2). If the community property is likely to be a community aggregated trait then exploring the link between microbial membership and community properties may lead to further understanding and perhaps an enhanced predictive power (Figure 2, Letter E). However, if the community property is likely an emergent property elucidating the microbial membership that contributes to the emergent property is, given the current understanding, unlikely to improve understanding of the drivers of that community property (Figure 2, Letter F). Formalizing microbiome research into a structured, conceptual framework should help the research community better focus on potential links between microbiome characteristics and system-level processes that are most likely to be detected empirically. This approach will also allow researchers working in different systems to test the same pathways among defined microbiome characteristics and thus increase the possibility of understanding the causal mechanism (or absence of causality) for observed correlations. Thus, future research endeavors will be most powerful if they focus on elucidating connections through the complete path of microbial ecology (Figure 2, blue arrow, Letters E, F, and G) and not direct connections between microbial membership or community properties and ecosystem processes (Figure 2, Letters I and J).

## Applying and Testing the Proposed Framework

Applying and testing the proposed framework will depend on the ability to more robustly characterize each category of microbial characteristics and to directly measure the arrows that connect each category (Figure 2). Both labeling/sorting approaches and phenotypic description of isolates provide an opportunity to better understand how microbial membership contributes to community properties (Figure 2, Letter E or F). Labeling and cell sorting approaches (e.g., fluorescent in situ hybridization (FISH) coupled with flow cytometry cell sorting,^35^ or immunocapture, such as with bromodeoxyuridine, BrdU)^36^ provide powerful tools to constrain the complexity of the microbiome and directly test hypotheses that link membership to community properties or microbial processes. For example, a study of an Arctic Ocean bacterial community labeled the actively growing component of the community using BrdU and then separated those populations from the rest of the community using an immunocapture technique to better understand the portion of the microbiome that was contributing to secondary production.^36^ In addition, physiological studies of isolates that are representative of important community properties have the potential to advance understanding of the role of phenotypic plasticity in structuring how constituent populations do or do not contribute to community properties (Figure 2, Letter E)^13^. Detailed studies of isolates of common environmental OTUs have demonstrated immense variation within a given OTU (i.e., “microdiversity”) that in part explains the challenge of linking membership to a community property^14^. For example, work on *Prochlorococcus* has led to a better understanding of how ecotypes within a single taxonomic unit (OTU) can lead to specialization in temperature, and substrate affinity^37^. OTUs that form a substantial portion of the microbiome’s sequence abundance provide potential candidates for further investigation of possible phenotypic plasticity and or microdiversity^14^. For example, a single phylotype of the class *Spartobacteria* within the phyla *Verrucomicrobia* was found to be present in a broad range of soil ecosystems and comprised as much as 31% of all 16S rRNA gene sequences returned from prairie soils^38^, making it an excellent candidate for targeted isolation and physiological studies. Studies of environmental isolates are essential in building a broader understanding of how community membership does or does not contribute to community properties (Figure 2, Letters E and F).

In addition to a better description of each category of microbial characteristics, an important step in moving from a correlative and descriptive approach to a causative and mechanistic approach comes in measuring the arrows represented by letters in Figure 2. There is a suite of powerful methods already being employed in microbial ecology that can actively measure many of the arrows illustrated in Figure 2, including: stable isotope probing of mixed communities^39^, single cell methods that can assay cells in the physiological state they occur in in the environment, and labeling individual cells with stable isotopes for single cell analyses^40^. Studies that use stable isotope probing or any form of tracking isotopically labeled substrates into a population have been successful in linking microbial membership to microbial processes (Figure 2, Letter H). For example, a study of a Scottish peatland used SIP to reveal that a single species of *Desulfosporosinus* was most likely responsible for the totality of sulfate reduction within the peatland even though it only comprised 0.006% of the retrieved 16S rRNA gene sequences^41^. In addition to this example of using SIP to link microbial membership and microbial processes (Figure 2, Letter H), there is a suite of cultivation independent techniques (such as Raman microspectroscopy (MS), NanoSIMS, or energy dispersive spectroscopy, EDS) that complement sequence-based microbiome analyses by reporting on the physiological and phenotypic characteristics of individual cells *in situ*. ^*40,42,43*^ Both Raman MS and NanoSims can be coupled with a range of *in situ* hybridization techniques (e.g., fluorescent in situ hybridization, FISH) to identify which populations are contributing to community properties (Figure 2, Letter E) or microbial processes (Figure 2, Letter H). For example, a study of a microbial consortia from the Sippewissett Salt Marsh on the coast of Massachusetts, USA used a combination of FISH and NanoSIMs to confirm a syntrophic association between a population of autotrophic purple sulfur bacteria and heterotrophic sulfate reducing bacteria (SRB)^44^. These existing methods of confirmatory ecophysiology allowed for direct measurements of the arrows connecting membership with microbial processes (in this case both carbon fixation and sulfate reduction, Figure 2, Letter H) in a stable microbial consortium.

These cultivation independent approaches also create the potential to begin to determine which community properties are emergent properties, and which are community aggregated traits. For example, microbial community biomass stoichiometry (e.g., biomass C:N or C:P) cannot currently be predicted (or even constrained) from a list of its constituent taxa (Figure 2, Letter F). However, microbial biomass stoichiometry is a community property with power to predict the microbial contribution to nutrient cycling (Figure 2, Letter G).^17^-^19^ Energy dispersive spectroscopy (EDS) has the power to measure the C:N:P of individual bacterial cells growing *in situ* (i.e. not in culture)^43^. The potential to couple EDS analysis with a phylogenetic label presents the opportunity to assay mixed microbial communities and assess the link between phylogenetic identity and biomass stoichiometry under natural conditions^45^. This approach would provide a direct link between community membership and a community property (e.g., biomass C:N, Figure 2, Letter E), that influences an important microbial process (i.e., nutrient recycling). These approaches applied in concert with sequence-based analyses have the potential to empirically link the categories of microbial information defined here (Figure 2) with the processes they influence, moving microbiome science from a descriptive and correlative approach to a mechanistic and causative approach.

## Designing microbiome research

The framework presented here provides one approach to formalize inquiry across microbiome science and encourages empirical linkages between the presence of organisms in a system and the processes that characterize that system. Whereas we draw examples from environmental microbiomes and the ecosystems they inhabit, this structured approach has the potential to benefit the analysis of microbiomes associated with other systems such as host organisms and those of the built environment. As important as establishing causal links among microbial membership, community properties, microbial processes, and ecosystem processes, is determining when these links are unlikely to be present. Research that indiscriminately seeks to identify correlations, which places all microbial characteristics on an equal plane and does not explicitly recognize the relationships between microbial characteristics, are likely to continue to yield conflicting and ambiguous results that not only fail to provide new insight into ecosystem processes, but also blur the connections that do exist. We suggest that rather than looking for linkages among microbiome membership and system-level processes in every study, research efforts would benefit from strategically targeting the linkages and processes for which an *a priori* understanding of microbial physiology should allow us to improve our understanding of the ecosystem process.

## Acknowledgements

This work is a product of the Next Generation of Ecosystem Indicators Working Group, supported by the USGS John Wesley Powell Center for Synthesis and Analysis. Preparation of this manuscript was supported by NSF DEB IOS #1456959 awarded to EKH. Chuck Pepe-Ranney and Ariane Peralta provided valuable feedback on previous versions of this manuscript. This paper is dedicated to Diana Nemergut, an integral part of our working group who passed away during the preparation of this manuscript. She is one of a kind.

Contribution
All listed authors have contributed to the conceptualization, writing, and preparation of the current manuscript.

